# Linking habitat and population dynamics to inform conservation benchmarks for data-limited salmon stocks

**DOI:** 10.1101/2021.03.22.436497

**Authors:** William I. Atlas, Carrie A. Holt, Daniel T. Selbie, Brendan M. Connors, Steve Cox-Rogers, Charmaine Carr-Harris, Eric Hertz, Jonathan W. Moore

## Abstract

Management of data-limited populations is a key challenge to the sustainability of fisheries around the world. For example, sockeye salmon (*Oncorhynchus nerka*) spawn and rear in many remote coastal watersheds of British Columbia (BC), Canada, making population assessment a challenge. Estimating conservation and management targets for these populations is particularly relevant given their importance to First Nations and commercial fisheries. Most sockeye salmon have obligate lake-rearing as juveniles, and total abundance is typically limited by production in rearing lakes. Although methods have been developed to estimate population capacity based on nursery lake photosynthetic rate (PR) and lake area or volume, they have not yet been widely incorporated into stock-recruit analyses. We tested the value of combining lake-based capacity estimates with traditional stock-recruit based approaches to assess population status using a hierarchical-Bayesian stock-recruit model for 70 populations across coastal BC. This analysis revealed regional variation in sockeye population productivity (Ricker α), with coastal stocks exhibiting lower mean productivity than those in interior watersheds. Using moderately-informative PR estimates of capacity as priors reduced model uncertainty, with a more than five-fold reduction in credible interval width for estimates of conservation benchmarks (e.g. S_MAX_ - spawner abundance at carrying capacity). We estimated that almost half of these remote sockeye stocks are below one commonly applied conservation benchmarks (S_MSY_), despite substantial reductions in fishing pressure in recent decades. Thus, habitat-based capacity estimates can dramatically reduce scientific uncertainty in model estimates of management targets that underpin sustainable sockeye fisheries. More generally, our analysis reveals opportunities to integrate spatial analyses of habitat characteristics with population models to inform conservation and management of exploited species where population data are limited.

## Introduction

Assessing population status and estimating conservation or management targets for data-limited fish populations is a major challenge to the sustainability of fisheries globally. In the absence of information to support management, many small unassessed fisheries around the world are depressed due to overfishing (Costello et al. 2012). This overfishing has resulted in the loss of billions of tonnes in potential fisheries yields per year (Ding et al. 2017), creating economic hardship and undermining food security in coastal communities (Golden et al. 2016). Given this challenge, a variety of approaches have been proposed for setting management targets without lengthy timeseries of population abundance and thus limited data. These range from management that relies on local knowledge rather than stock-assessment data (Johannes 1998), to quantitative approaches such as meta-analysis that combine information from multiple sources to reduce uncertainty associated with sparse timeseries of abundance within a single population (Myers and Mertz 1998, Punt et al. 2011).

Meta-analytic approaches allow researchers to combine insights from multiple populations and are a valuable tool for understanding and managing populations with limited data (Myers and Mertz 1998). These analyses rely on the assumption that population parameters are drawn from a shared underlying distribution. Under this assumption, population parameters – for example the maximum annual reproductive rate (alpha) in a Ricker stock-recruit model (Ricker 1954) – can be readily estimated for data-poor populations, as the model can borrow information from populations with more robust time series (Gelman 2006, Thorson and Minto 2015). Similarly, researchers now routinely pool information across populations using hierarchical-Bayesian methods which assume that some population parameters are drawn from common hyper-distributions (Punt and Hilborn 1997). These approaches have been used by many researchers to examine stock-recruit relationships (e.g. Liermann and Hilborn 1997, Michielsens and McAllister 2004) and understand the impacts of climate on recruitment across multiple populations (e.g. Mueter et al. 2002, Malick et al. 2016).

Another potential approach to inform management in data-limited fisheries involves predicting management-relevant parameters based on habitat information, when habitat limits production (e.g. Sundblad et al. 2014). For fish populations where density-dependent dynamics occur in well-delineated habitats, such as the use of freshwater habitat by juvenile salmon, habitat quantity and food web productivity can impose constraints on the carrying capacity of fish populations. For example, the amount and gradient of available stream-rearing habitat has been used to predict coho production (Bradford et al. 1997, Bocking and Peacock 2004), and accessible watershed area has been used to inform estimation of population parameters in data-limited Chinook populations (Parken et al. 2006, Liermann et al. 2011). While stock-recruit modeling is data intensive, often requiring decades-long time series, these habitat-based models offer the advantage of only requiring information on the amount or quality of available habitat that could be estimated remotely using geospatial analysis, or with as little as a single year of field sampling. By coupling data from populations with intensive population monitoring and the known habitat constraints for the species of interest, researchers can model the underlying relationship between habitat conditions and population parameters estimated from stock-recruit timeseries (e.g. Hume et al. 1996, Parken et al. 2006). This relationship can then be extended to estimate management targets such as carrying capacity or maximum sustainable yield (MSY) for fish populations without long timeseries of stock-recruit data. These habitat-based estimates of capacity can be merged with stock-recruit analyses in a Bayesian framework, either through their inclusion in the model as a covariate modifying the strength of density dependence (e.g. Liermann et al. 2010), or as a population-specific prior on the spawner abundance at the produces maximum recruitment (e.g. Korman et al. 2013).

Sockeye salmon are a semelparous and anadromous species of Pacific Salmon, and are a primary target of commercial, recreational, and First Nations subsistence fisheries in coastal British Columbia and Alaska. Sockeye salmon generally have an obligate juvenile lake-rearing phase of one or two years prior to their seaward migration, during which time they feed on zooplankton and invertebrates (Groot and Margolis 1991). Given this dependence on rearing habitat in lakes, lake size and food-web productivity can control the carrying capacity of sockeye populations (Juday et al. 1932, Hyatt and Stockner 1985, Shortreed et al. 2001). In recent decades, researchers and managers in Alaska and British Columbia have developed different rearing-capacity models for sockeye bearing lakes, predicated upon these physical and ecological constraints. Among these models is the euphotic volume model which relates sockeye rearing capacity to the amount of volume in a lake’s euphotic zone, as surrogates for lake productivity (Koenings and Burkett 1987). However, models which measure lake productivity directly tend to produce more reliable predictions of fish production (Downing 1990), and Hume et al. (1996) developed a model that related lake photosynthetic rate (PR) to sockeye production. The PR model scales-up monthly estimates of photosynthetic rates to total annual growing season carbon production, using lake area and a defined growing season length. Hume et al. (1996) used data from several populations with juvenile population enumeration (i.e. fall fry or smolt) and lake PR monitoring to model the empirical relationship between smolt output and total autotrophic production. This relationship between annual PR and smolt output has subsequently been used to predict carrying capacity for approximately 60 lakes in coastal British Columbia (e.g. Shortreed et al. 1998, Shortreed et al. 2001, Shortreed et al. 2007). To date, these data have not been fully integrated with existing timeseries of spawner abundance to estimate sustainable harvest rates or evaluate conservation status of sockeye populations, an important step towards providing management advice particularly in populations where stock-recruit data are scarce (e.g. Cox-Rogers 2010).

Canada’s Wild Salmon Policy (WSP) calls for the establishment of conservation benchmarks for evaluating population status and implementing management and recovery efforts. While sockeye salmon are relatively well-studied in many parts of their range, timeseries of spawner abundance and recruitment are often sparse in more remote regions and for smaller populations, creating challenges for setting management goals and conservation benchmarks. Conservation benchmarks which rely solely on biological information differ from management targets which consider socio-economic factors (Holt and Irvine 2013). The WSP was originally adopted in 2005 by Fisheries and Oceans Canada (DFO), with the goal of safeguarding the genetic and ecological diversity of wild Pacific Salmon for the benefit of Canadians in perpetuity (DFO 2005). Recent updates to the Fisheries Act also include provisions requiring management for sustainable fisheries, with mandatory rebuilding plans when populations drop below biological reference points. However, the implementation of management and conservation actions for these populations as mandated by the WSP and Fisheries Act, is currently hindered by data-limitations for many populations.

The north and central coast (NCC) of BC is a region where the development of such benchmarks has proven a challenge but are of timely importance. The NCC is remote and vast, covering more than 175,000 km^2^, and there are more than 120 genetically and demographically distinct populations of lake-type sockeye salmon, designated as Conservation Units (CU) under the WSP. A few well-studied populations have been assessed to be depressed (McKinnell et al. 2001; Connors et al. 2019; Price et al. 2019), but most watersheds in the NCC region are remote, and only accessible by boat or air, making consistent population monitoring logistically challenging and costly. Further, many of these populations are small, with average run sizes of less than 10,000 fish, and monitoring efforts have historically been focused on the largest and most commercially important populations. Understanding the status and capacity of remote coastal sockeye salmon populations is particularly important to coastal First Nations. These communities are increasingly assuming leadership of resource monitoring and management within their traditional territories, and managing food, social and ceremonial sockeye fisheries for sustainable economic and cultural benefits is a primary goal (e.g. Atlas et al. 2017). Thus, methods that integrate multiple sources of information on populations and their habitats are needed to inform management of data-limited sockeye salmon populations.

Here we integrate habitat-based and population-dynamic based approaches to inform conservation and management for 70 populations of sockeye salmon in coastal BC. Specifically, we developed a hierarchical-Bayesian Ricker model of spawner-recruit dynamics for sockeye salmon, integrating information on lake productivity and size through the inclusion of habitat-based estimates of carrying capacity as prior information, and asked the following questions: (1) how does the inclusion of a habitat information affect estimates of population productivity and capacity and (2) how do current sockeye population abundances in the NCC compare to conservation benchmarks derived using habitat-based priors?

## Methods

### Overview

We estimated the following parameters for each of the 70 sockeye populations: 1) population abundance over the last 15 years, 2) spawning abundance at carrying capacity (S_MAX_), and 3) spawning abundance at maximum sustainable yield (S_MSY_). These parameters can inform conservation benchmarks and fishery management. For each of 70 sockeye populations, we fit stock-recruitment models to timeseries of spawner abundance, catch, and average brood year age composition (English et al. 2016). These stock-recruitment models incorporated habitat-based priors drawn from more than 20 years of limnological assessments conducted by DFO’s Lakes Research Program in sockeye-bearing lakes of the NCC (e.g. Shortreed et al. 1998, Shortreed et al. 2001, Shortreed et al. 2007). These data were then combined in a series of models to evaluate the degree to which information on the productivity of rearing habitats was a suitable source of prior information on population carrying capacity, and how the inclusion of these priors affected estimates of carrying capacity (S_MAX_) and spawner abundance at maximum sustainable yield (S_MSY_).

### Photosynthetic rate model

We used published estimates of lake carrying capacity (S_MAX___PR_) derived from the photosynthetic rate (PR) model first developed by Hume et al. (1996) and refined by Shortreed et al. (2000) (Table S1). Hourly and daily photosynthetic rates were estimated *in situ* using light and dark bottle incubations within the euphotic zone, which measure autotrophic uptake of inoculated ^14^C isotopes in relation to incident light levels (see Shortreed et al. 1998 for detailed methods). These hourly estimates of PR were typically made multiples times over a growing season. They were then temporally expanded to daily rates and seasonal mean photosynthetic rates based upon growing season length (May 1^st^ - October 31^st^) and lake surface area, to estimate total annual growing season carbon production (Shortreed et al. 2000).

The PR model assumes that sockeye populations are limited by lake productivity and area, and previous research in BC has suggested that in most cases this assumption is valid (Shortreed et al. 2001). Drawing on the work of Koenings and Burkett (1987), Hume et al. (1996) used the correlation between empirically-derived estimates of spawner abundance at carrying capacity (S_MAX_) and total annual primary production (PR_TOTAL_), to estimate S_MAX_ for lakes using only PR data (S_MAX___PR_). This effort yielded a relationship between lake productivity and sockeye population capacity which has subsequently been used to estimate carrying capacity in lakes across the NCC (Shortreed et al. 1998, Shortreed et al. 2001, Shortreed et al. 2007). The model assumes a fixed relationship between lake productivity and maximum smolt output, thus PR_TOTAL_ (tons·C·year^-1^) can be multiplied by the constant 187 (spawners· tons·C^-1^) to yield an estimate of the number of adult spawners required to maximize smolt production (Shortreed et al. 2000) (Equation 1).

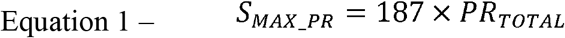

In most applications, PR measurements are made throughout the growing season, accounting for seasonality in primary production (see Shortreed et al. 2001 for summary). However, across the numerous remote NCC lakes access is difficult and assessment costs are high. Thus, single estimates of PR made in late-August or early-September have been successfully related to seasonal mean PR values yielding a correction factor of 0.748, which can be applied to produce estimates of seasonal mean PR (Cox-Rogers et al. 2004). When estimates of annual primary production (PR_TOTAL_) were available for multiple years, we used the average of these estimates as our prior. In other cases, multiple lakes support rearing sockeye within a single population (e.g. Elbow, Lonesome, and Rainbow Lakes for Atnarko sockeye), and priors reflected the sum of productivity of all lakes known to support sockeye rearing. For more information on data sources and methods see the supplemental materials (Table S1).

While field measurements of PR were only available for 40 of 70 populations in our study, recent parallel modeling efforts have led to the creation of a landscape-scale predictive model for lake PR across the NCC (Atlas et al. 2020). These efforts examined a series of potential landscape and geographic variables as predictors of S_MAX___PR_ and found there were strong regional trends and that sockeye lake productivity was well predicted by whether the lake is clear, humic stained or glacially turbid. We therefore used predicted S_MAX___PR_ values from this landscape model as priors for S_MAX_ in populations where rearing lakes were unsampled.

### Escapement and catch data

We used abundance and harvest data from 70 populations on the NCC collected since the 1950s by DFO. Following English et al. (2016), we assumed that neighboring populations within statistical areas have similar run timing and distribution and are therefore similarly vulnerable to coastal fisheries. So, in cases where harvest data were not available for a specific population, we used the average harvest rate for other populations in the same DFO statistical area to reconstruct catch. These data represent the best available information on harvest rates over time but should be interpreted with caution when neighboring populations diverge in their vulnerability to fisheries. Harvest rates ranged from more than 70% in the early part of the timeseries for some populations to less than 10% in more recent years. Using available age data (English et al. 2016) as well as model estimates of age composition (see below), we constructed brood tables assigning recruits to previous parent cohorts to estimate the relationship between spawner abundance and recruitment. We examined each population timeseries individually, identifying and removing years which produced unreasonably high estimates of per capita recruitment (more than 20 adult recruits per spawner). These outliers were likely because of poor data quality or extrapolated estimates of harvest that inflated per capita recruitment. We also dropped data points of fewer than 100 spawners, as population sizes that small are rare and are likely a reflection of poor data quality. This process resulted in the elimination of 73 data points out of a possible 1,850 spawner-recruit pairs. Data richness varied widely across the 70 populations of interest, with the number of spawner-recruit pairs ranging from 4 to 57.

### Age structure

The quality and availability of age data was highly variable across populations. In some large and commercially-important populations (Atnarko, Babine, Long, Meziadin, and Owekino) estimates of annual age data are available. Specifically, for Babine, age-composition data is available throughout the timeseries. For Atnarko, age data is available in 33 years from 1976 to 2016 and in years missing age data we assumed age composition was equal to the long-term average. In the remaining three populations, annual age data was available only since 1989, so estimated recruitment to cohorts after 1986 reflects annual age variation, while earlier estimates of recruitment reflect mean age composition. In many other populations (n = 19), age estimates are limited to a few years, and we used average brood-year age composition values reported in English et al. (2016), data from Todd and Dickinson (1970) for Bowser Lake, and brood-year age composition reconstructed from scale and otoliths collected since 2012 during annual monitoring in Koeye, Namu, Port John, and Kadjusdis (W.Atlas unpublished data).

For the remaining 45 populations without age data we modelled the available multinomial age proportion data against environmental correlates using Dirichlet regression (R-package DirichletReg version 0.6-3; Maier 2015) and using model outputs to predict age structure for populations lacking age data. Salmon exhibit considerable intraspecific variation in life histories and age at maturity (Quinn 2005), and this variation is often linked to differences in climate or hydrology within their natal watersheds (Beechie et al. 2006). We found maximum watershed elevation was the best predictor of age composition; higher elevation watersheds tended to have older fish. This relationship was used to predict age structure for watersheds lacking data (Figure S1, Table S2). This novel approach to predicting population age composition across the landscape facilitated the creation of brood tables for stock-recruit analysis, however these estimates of population age structure are uncertain. Accordingly, estimates of S_MAX_ and S_MSY_ for populations lacking age data should be interpreted with greater caution.

### Stock-recruit dynamics and alternative modeling approaches

We modeled density-dependent population dynamics for each timeseries of spawner abundance (S) and recruitment (R) using a Ricker model (Ricker 1954), where α controls the per-capita productivity (slope) at the origin, and β dictates the strength of density-dependence. This equation is widely used in part because it can be adapted to a linear relationship by taking the natural-log value of the number of recruits per spawner at a given population size. Spawner abundance at carrying capacity (S_MAX_) is then estimated as the reciprocal of β. To ease interpretation and integration of data from multiple data sources, priors for β_*i*_ were specified in relation to S_MAX*i*_. For each population α was assumed to be lognormally distributed with a mean of 1, and a standard deviation of 1, creating a moderately informative prior that bounded percapita productivity within biologically plausible values (Equation 2).

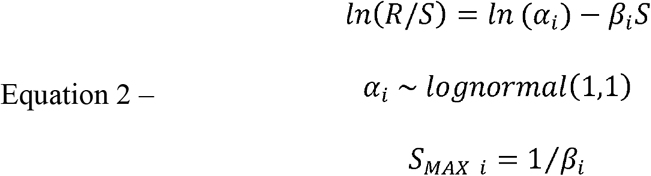

To evaluate the effects of estimating productivity (*α*) hierarchically and using S_MAX___PR_ as a prior, we estimated Ricker stock-recruit parameters for each population (*i*) using four alternative models:

1. Productivity (*α*) values were estimated independently for each of the 70 populations, with a uniform prior for S_MAX*i*_ ranging from 0 to 10 times the maximum observed escapement in a population (*MaxE_i_*) (Equation 3).

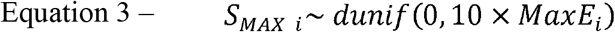
2. Productivity values were estimated hierarchically with vales of α drawn from a hyper-distribution with a mean of *mu.α* and a precision of *tau.α* (Equation 4), and a uniform prior on S_MAX*i*_ (Equation 3).We used uninformative uniform priors for *mu.α* and *tau.a*. The sockeye bearing watersheds NCC span broad hydrological and climatological gradients, from low elevation coastal bog-forest through deep mountainous fjords, and into the interior plateau. Given the potential differences in environmental conditions influencing sockeye productivity across these three regions, we evaluated whether there was support for including three regional priors on *mu.α*, by fitting model 2 with and without region-specific hyper-distributions that corresponded to the biogeoclimatic zones defined by Holtby and Ciruna (2007) for Pacific salmon (coastal, fjord, interior). We then compared the density distributions of the regional *mu.α* estimates to evaluate the degree of statistical support for regional differences in productivity.

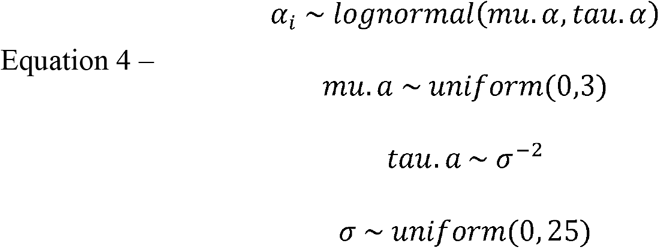
3. We used a semi-informative log-normal prior for S_MAX*i*_, based on the estimated productive capacity of the rearing lake. This prior was given a mean of S_MAX___PR*i*_ (Table S1) and a standard deviation of 0.9, yielding a moderately informative prior with values typically spanning a range of approximately 0.1 to 10 times the long-term mean population size and the highest probability density at S_MAX___PR*i*_ (Equation 5). Productivity (*α*) values were estimated independently for each population as in model 1 (Equation 2).

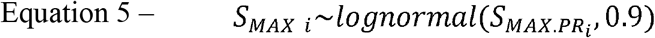
4. Alpha values were estimated hierarchically as in model 2 (Equation 4), with a semi-informative prior based on S_MAX___PR_ as in model 3 (Equation 5).

Models were then run for 100,000 iterations in JAGS using the statistical program R, with three parallel chains, and a burn in period of 50,000 iterations. Model convergence was evaluated visually using trace plots.

For all models we estimated population size at maximum sustained yield (S_MSY_) using Scheuerell’s method, where *W* is the solution to Lambert’s function, implemented in the R-package gsl (Scheuerell 2016) (Equation 6). Posterior density distributions of S_MSY_ for each population were estimated using 10,000 randomly sampled α and β values from the MCMC, and mean and confidence intervals were estimated using the R-package HDInterval (Meredith and Krushke 2020).

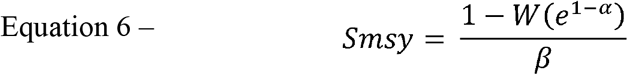

### Validation of PR model as S_MAX_prior

Given the variability in the availability and quality of data across the NCC, populations were differentiated into two groups: (1) those with relatively high-quality stock-recruit and age data (n = 12), and (2) those with lower quality stock-recruit timeseries and poor or missing age data where stock-recruit modeling may produce biased or uncertain estimates of population parameters in the absence of informative priors (n = 58). We compared estimates of spawner abundance at carrying capacity derived from the PR model (S_MAX___PR_) (Equation 1), and those estimated from the Ricker stock-recruit model (S_MAX_) (Equation 2), for populations with more than 25 stock-recruit pairs and where there was available information on population age structure (group one). In this exercise we fit model 1 - with independent estimates of productivity (*α*) and uninformative uniform priors on spawner abundance at carrying capacity (S_MAX_) - to stock-recruit timeseries from 12 populations which met our standards for data quality (Alastair, Babine, Bella Coola, Kitsumkalum, Koeye, Lakelse, Long, Meziadin, Morrison, Namu, Owikeno and Tankeeah).

We then fit a linear regression with the intercept constrained through the origin, to compare estimates of spawner abundance at carrying capacity values from the stock-recruit and PR model. We considered a slope that did not statistically differ from 1 as support for using estimates of S_MAX___PR_ as a prior for Ricker S_MAX_ in subsequent stock-recruit modeling.

### Evaluation of model performance

To evaluate the benefit of including habitat-based priors on S_MAX_ we standardized model estimates of S_MAX_ by the mean timeseries abundance for each population and compared the width of credible intervals (CI) between models with (Model 4) and without PR priors (Model 2). For the purpose of this analysis we removed five populations – Bear, Kitwanga, Morice, Motase, and Swan – which are not thought to be limited by lake rearing habitat (Shortreed et al. 2001, Cox-Rogers 2010).

### Assessment of population status and regional productivity trends

To facilitate comparisons across populations spanning several orders of magnitude in average abundance, median estimates of spawner abundance at carrying capacity (S_MAX_) as well as lower and upper 95% credible intervals were standardized by mean escapement across the timeseries. A lack of recent spawner abundance data for many populations limited our ability to evaluate recent escapement trends, so we compared mean escapement (S) since 2000 to modeled estimates of S_MAX_ and S_MSY_ to assesses the relative population status of each population. Abundance relative to S_MSY_ has previously been proposed as a benchmark for delineating populations that are considered healthy and those where reductions in abundance pose the risk of further population declines and erosion of fisheries opportunities (Holt and Bradford 2011; Costello et al. 2012). Therefore, for the purpose of our analysis populations where S/S_MSY_ >1 are above conservation benchmarks, populations where S/S_MSY_<1 are currently below conservation benchmarks. Populations with fewer than 3 estimates of escapement since 2000 were dropped from the analysis, leaving a total of 54 populations where status could be assessed.

## Results

Across all regions, hierarchical modeling of population productivity (*α*) resulted in more constrained estimates of productivity, particularly in the most data-limited populations (Figure 2). On average, 95% CIs of *α* for the single population model (Model 1) spanned 2.79 R/S, while hierarchical estimation of *α* reduced the mean 95% CI to 2.17 R/S (Model 2). Hierarchical estimates of productivity (*α*) were shifted towards the mean of the hyper-distribution, resulting in a reduction in the range of values falling within the 95% credible intervals for *α* for all but four populations. There was statistical support for modeling productivity (*α*) hierarchically by region. Different regions had different productivities—specifically, hierarchical mean productivity (*mu.α*) of low elevation coastal sockeye stocks was substantially lower (2.78 R/S; 95% CI 2.72 – 2.98) than interior sockeye (3.76 R/S; 95% CI 3.09 – 4.69), with non-overlapping confidence intervals. Hierarchical mean productivity for coastal fjord populations was intermediate between low coastal and interior regions (*mu.α* = 3.08 R/S; 95% CI 2.74 – 3.91) (Figure 2). Given this statistical evidence for regional differences we used the regional-hierarchical productivity model as the base model for comparing estimates of carrying capacity and S_MSY_ with uniform and lake PR-based priors (Models 2 & 4). Across the 70 populations we analyzed, mean productivity in the single population and regional-hierarchical model were 3.13 and 3.09 (R/S) respectively (Table S4). Estimated *α* values in populations with the lowest and highest productivities experienced the greatest shrinkage when estimated hierarchically, with the lowest median *α* value increasing from 1.40 to 1.99 and the highest median *α* declining from 8.90 to 6.31 between our single population (Model 1) and regional-hierarchical models (Model 2).

**Figure 1:**
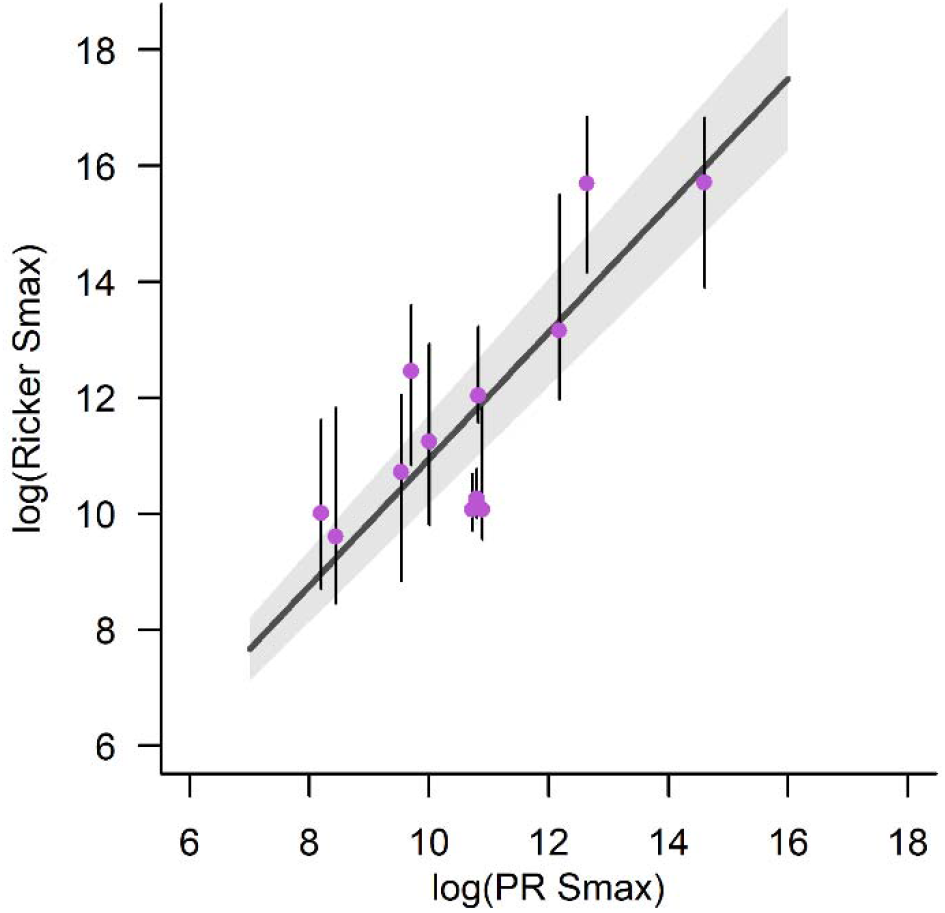
Comparison of stock-recruit derived estimates of spawner abundance at capacity to S_MAX___PR_ for 12 data-rich lakes.

**Figure 2:**
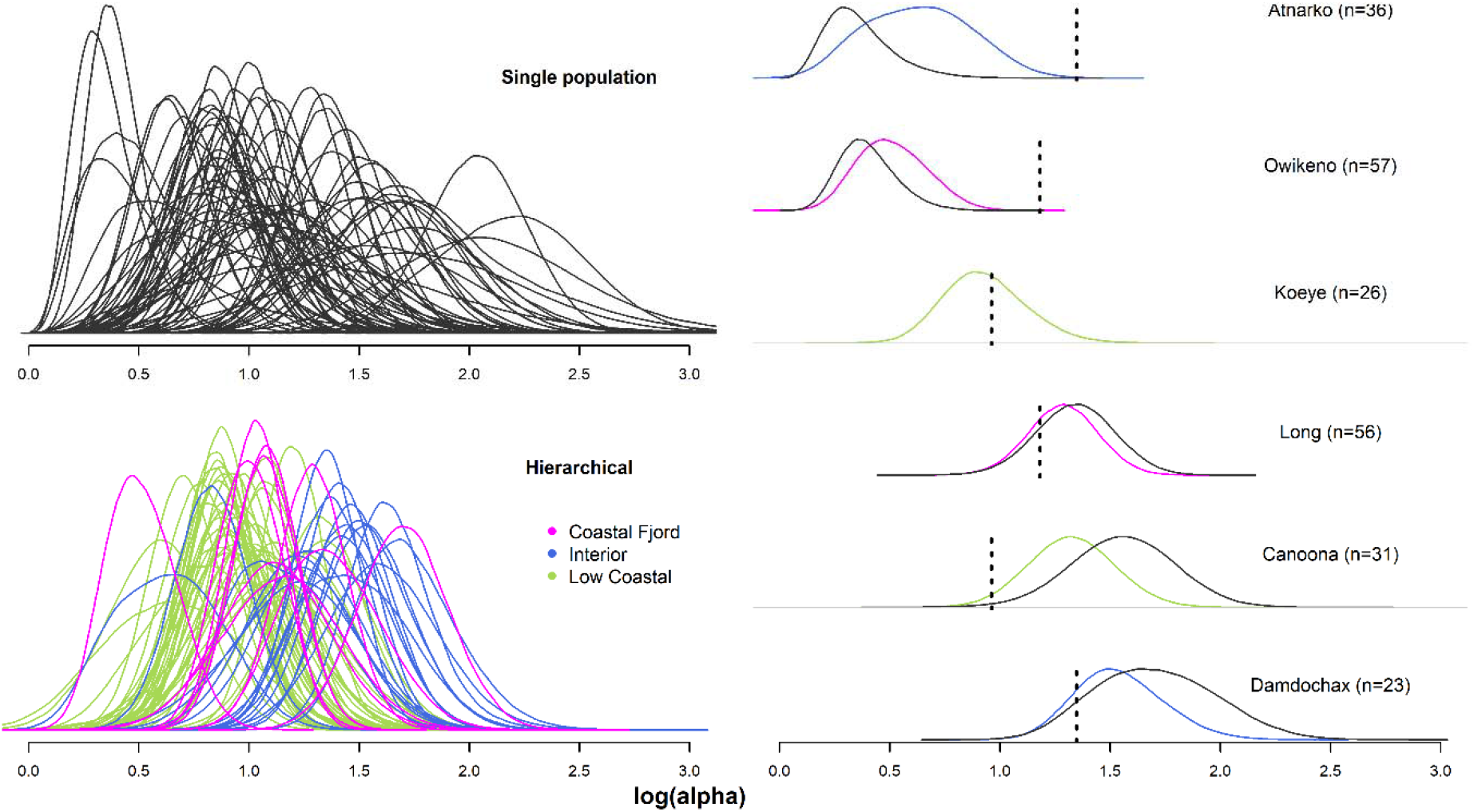
Estimated values for the natural log of alpha (population productivity) for all 70 populations with estimates from the independent (non-hiearchical) population model shown in black and estimates from the regional-hierarchical model shown in pink (Coastal Fjord), green (Low Coastal), and blue (Interior). Individual population plots show shrinkage resulting from hierarchical estimation of alpha, with the vertical dashed line representing the hierarchical mean alpha for a given region.

Given the observed shrinkage in *α* towards the hierarchical mean value, hierarchical modeling of *α* in our model with habitat-based priors (Model 4) resulted in subtle but important changes in estimates of S_MSY_ relative to the same model fit with independent *α* for each stock (Model 3). While median estimates of S_MSY_ did not differ appreciably between the two models (< 3% mean difference), fitting *α* hierarchically reduced the 95% credible intervals for S_MSY_ by an average of 4.5%. In particular, estimates of S_MSY_ for populations with the lowest and highest productivity (*α*) had the greatest discrepancy in median estimates and uncertainty. For example, the median estimates S_MSY_ for Owikeno Lake sockeye – the population with the lowest estimated productivity – were 448,953 and 445,947 spawners respectively for the independent and hierarchical models. But the 95% credible interval for estimates of S_MSY_ for the model with independent estimates of *α* ranged from 275,780 to 731,941 spawners, while the hierarchical model estimates ranged from 303,913 to 649,841 spawners. By contrast, median estimates of S_MSY_ for Fred Wright Lake – the population with the highest estimated productivity – were much lower when productivity was estimated independently (9,743 spawners CI: 5,727 to 18,794) than when it was estimated hierarchically (11,610 spawners, CI: 6,189 to 25,632).

Our initial analysis of 12 data-rich systems supported the use of S_MAX___PR_ as prior information for stock-recruit derived estimates of S_MAX_. The linear relationship between Ricker estimates of S_MAX_ and S_MAX___PR_ had an estimated slope of 1.05 (95% CI 0.99 – 1.11) (Figure 1). Thus, there was agreement between estimates of sockeye carrying capacity based on habitat and limnology, and estimates based on relatively high quality timeseries of abundance and age structure. We therefore used both empirical and predicted PR-based estimates of capacity (S_MAX___PR_) as priors on S_MAX_ for all 70 populations.

The inclusion of lake-based priors for S_MAX_ tended to reduce uncertainty in estimated population parameters. In many populations, model fits for the hierarchical Ricker model with uninformative priors produced highly uncertain estimates of S_MAX_ and S_MSY_ (Model 2). In the absence of prior information on lake capacity (S_MAX___PR_), only 17 populations had 95% credible intervals for S_MAX_ that spanned less than two times their mean escapement, and the median credible interval for S_MAX_ spanned a range of abundances that were 5 times the mean escapement. In some populations, parameter estimates were extremely uncertain with credible intervals for S_MAX_ in 17 populations that were greater than 20 times their mean escapement, and a maximum of almost 48 times mean escapement (Figure 3, Table S4). Informative priors were particularly valuable in populations with high uncertainty in Ricker model fits (Figure 4). For 63 of the 65 populations (excluding aforementioned populations not limited by lake productivity), including informative priors (Model 4) resulted in greater certainty in estimates of S_MAX_, with a median 95% CI spanning 1.58 times the mean escapement (Figure 3, Table S4). With the use of informative priors, the number of populations for which the range of S_MAX_ estimates fell within two times mean escapement increased to 36 of 65 (from 17 of 65 with uninformative priors). There was not always agreement between values of capacity estimated from stock-recruit models (S_MAX_) and those estimated only from lake productivity (S_MAX___PR_), which may highlight populations where lake productivity is not the primary factor limiting sockeye production or non-stationarity in recruitment dynamics.

**Figure 3:**
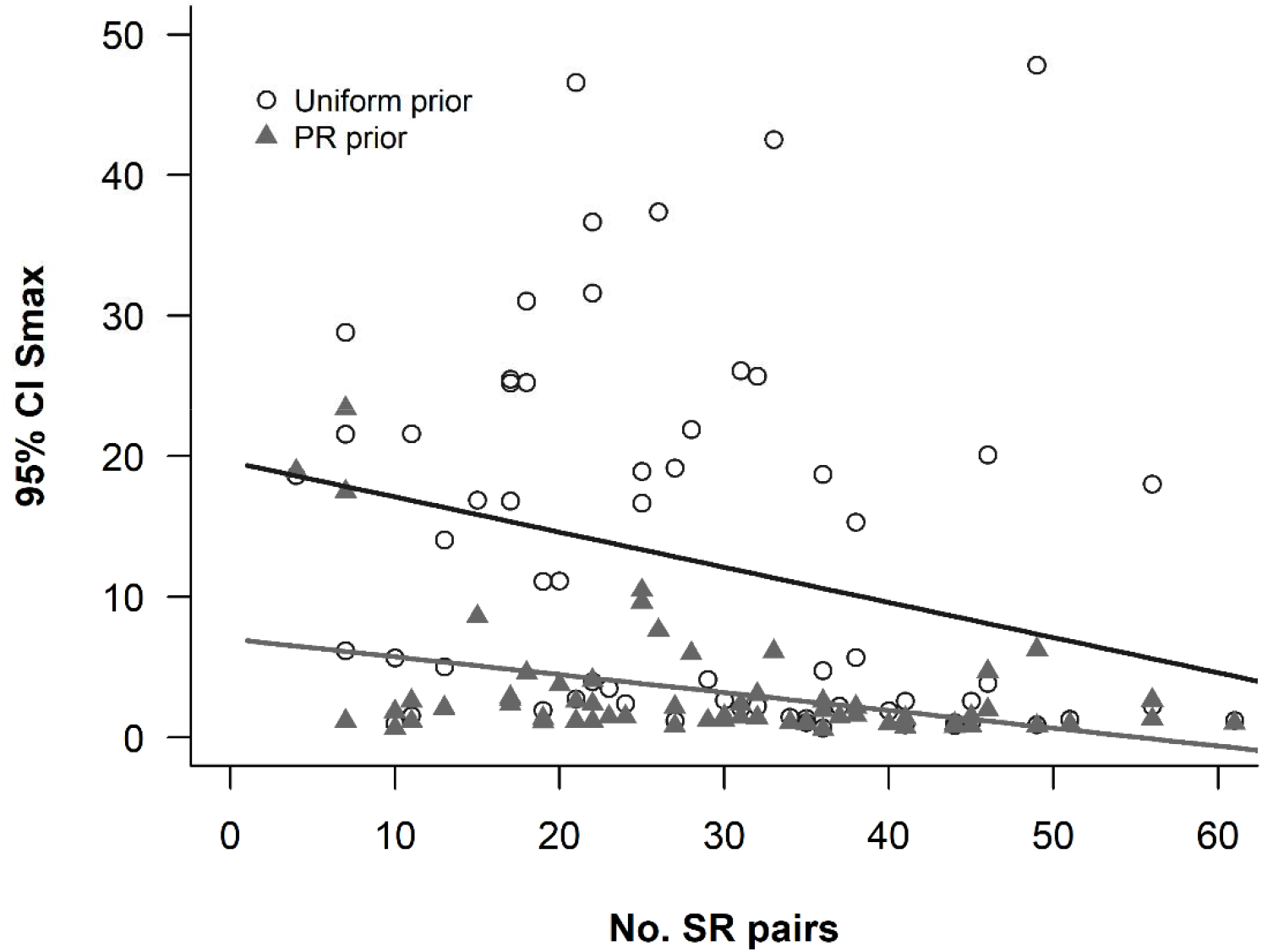
Relationship between width of 95% credible interval for S_MAX_ and the number of stock recruit pairs used in modeling, for models with uninformative uniform priors (open circles), and lake PR based priors (solid triangles).

**Figure 4:**
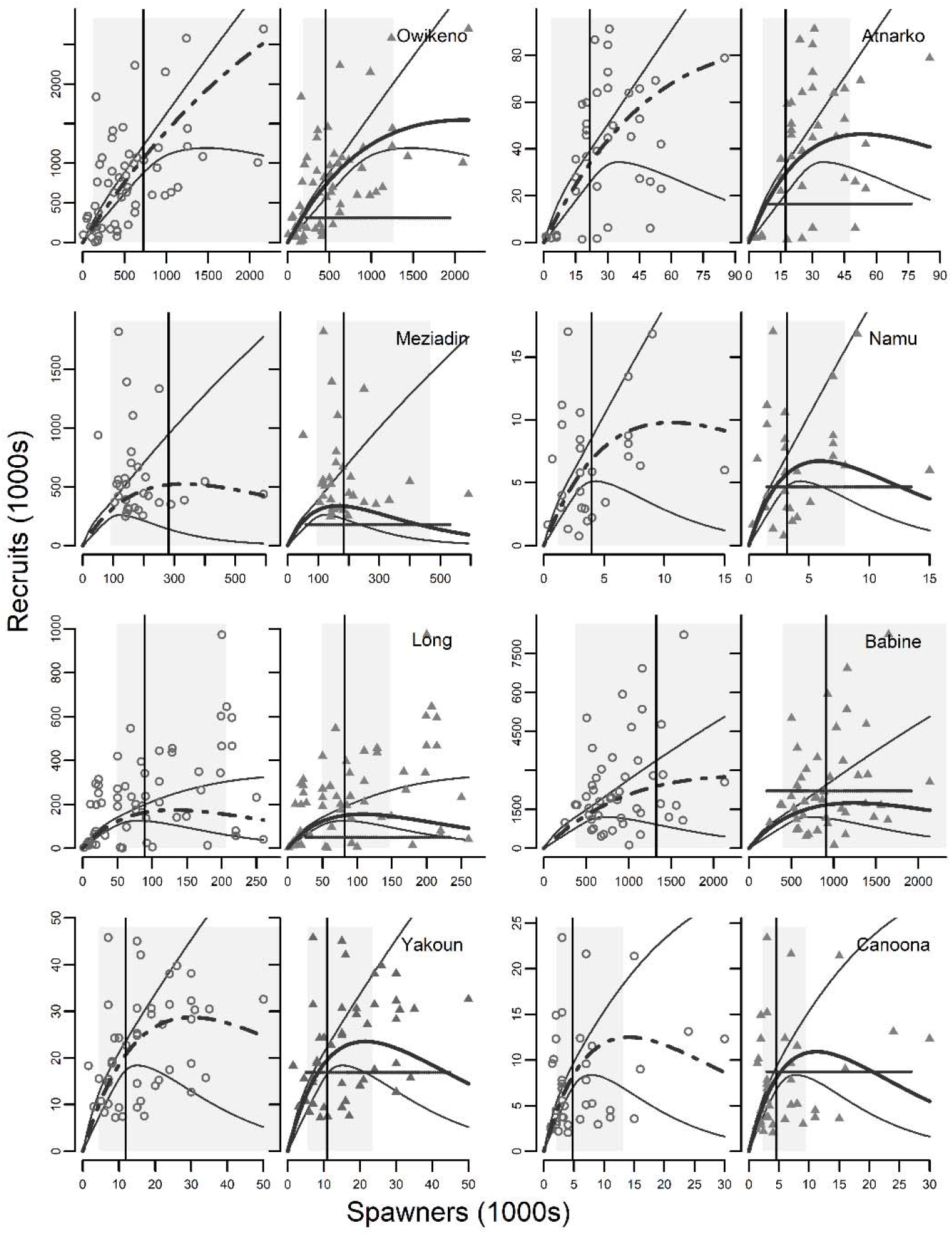
Stock recruit relationships with uniform (circles and dashed lines) and lake PR-based priors (triangles and solid lines) on S_MAX_ for eight of the 70 populations on the NCC with spawner escapement data. Gray rectangles and vertical lines are 95% credible interval and median S_MSY_. Horizontal dashed lines indicate PR priors on spawner abundance at carrying capacity (S_MAX___PR_).

### Population status

While the lack of recent escapement data made assessment impossible for some systems (16 of 70 populations), we evaluated population status relative to S_MSY_ and S_MAX_ for the remaining 56 systems with three or more years data since 2000. About half of sockeye populations on the NCC have experienced recent spawner abundances above those predicted to maximize fisheries yield (S_MSY_). For models (2) with uniform and (4) lake-based priors, recent sockeye escapement averaged 1.16 and 1.04 times the median estimates of S_MSY_ and 0.58 and 0.5 of S_MAX_ respectively. Thus, the current average population abundance is approximately half the long-term carrying capacity, and at or near levels of abundance that maximize sustainable yield even though harvest rates have been curtailed dramatically.

Given differences in estimates of S_MSY_ between models with (Model 4) and without habitat-based priors (Model 2) on capacity, the two models produced slightly different assessments of status among some populations. The uniform prior model (Model 2) estimated that 25 of 56 populations had recent mean escapements above their median S_MSY_ (Figure 5, circles). When lake-based priors for S_MAX_ were included (Model 4), 30 populations were above median S_MSY_ (Figure 5, triangles). Given uncertainty in estimates of S_MSY_, 14 populations had recent escapements which did not overlap the 95% CI of S_MSY_ when we included prior information on lake capacity. In our uniform prior model, estimates of S_MSY_ were more uncertain and 13 populations did not fall within the 95% CI for S_MSY_. There was consensus between models that 24 sockeye populations had a greater than 50% probability of being below S_MSY_, suggesting that recent escapements in these populations are depressed relative to both their habitat potential, and previously observed patterns of productivity. (Swan, Mary Cove, Kitkiata, Fred Wright, Kadjusdis, Kwakwa, Damdochax, Owikeno, Asitika, Kitwancool, Kitlope, Atnarko, Long, Port John, Meziadin, Namu, Curtis Inlet, Bloomfield, Yakoun, Morice, Marian Eden, Price, Skidegate, and Freeda Lake)

**Figure 5:**
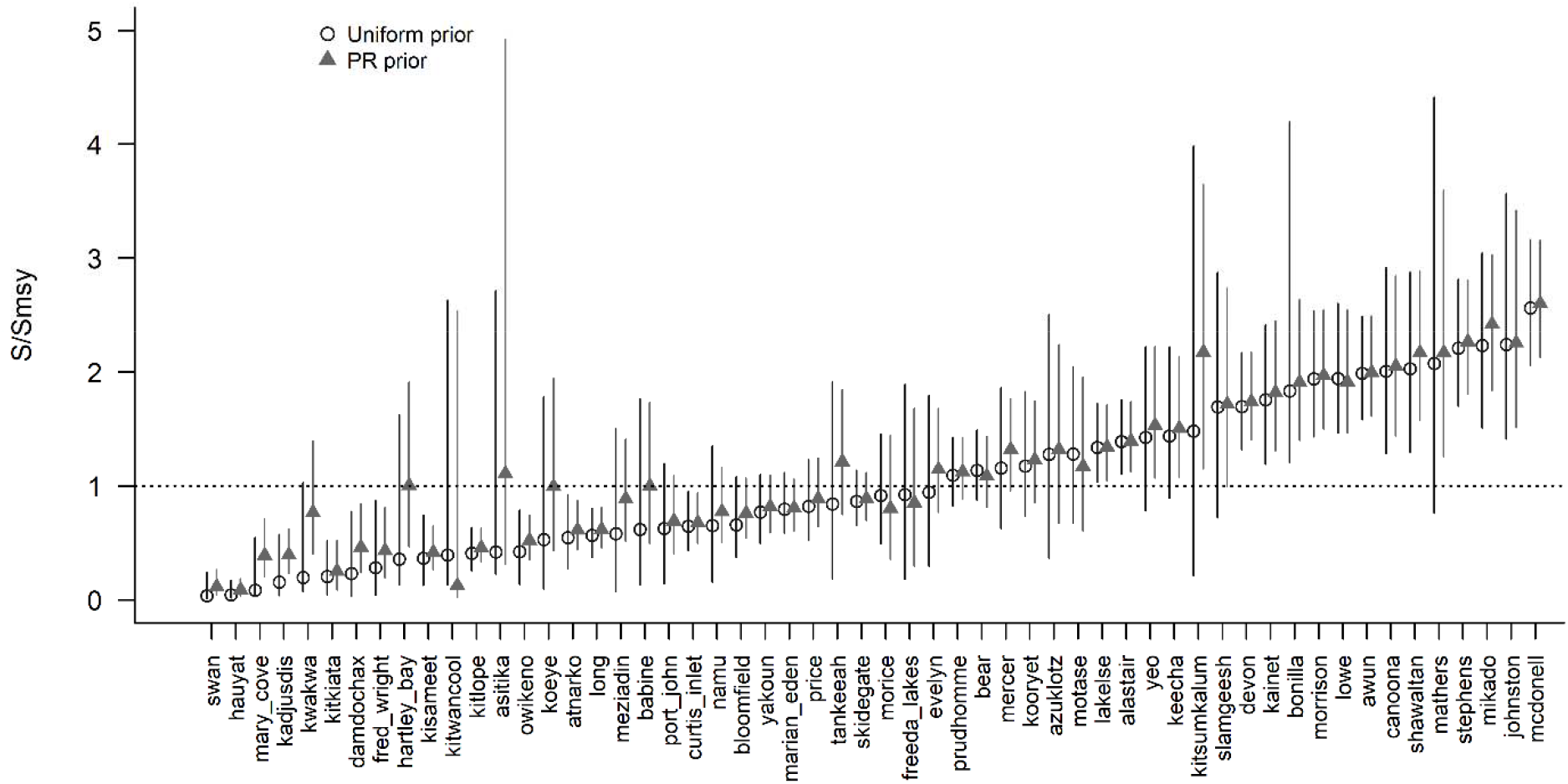
Relationship between recent escapements (S) and estimated spawner abundance at MSY (S_MSY_) for 54 populations with recent escapement data. Vertical lines represent 95% credible intervals for estimated stock status. The dashed line is indicative of a population size equal to MSY.

## Discussion

Here we developed and applied stock recruitment models that incorporate habitat information to assess the conservation status of data deficient sockeye salmon populations. We used a hierarchical-Bayesian Ricker model that combines spawner-recruit dynamics and lake habitat information to estimate population productivity (*α*), spawner abundance at carrying capacity (S_MAX_), and the escapement levels predicted to maximize fisheries yield (S_MSY_) for 70 populations of sockeye across the North and Central Coast (NCC) of British Columbia. Comparing estimates of S_MAX_ derived from stock-recruit methods with lake productivity-based capacity estimates (S_MAX___PR_) from 12 populations with high quality data indicated that the two sources of information are comparable, supporting the use of lake-based photosynthetic rate (PR) model estimates as a prior for S_MAX_ in stock-recruit models. For many of these populations, our analyses represent the first effort to estimate stock-recruit parameters necessary for conservation and management planning, and extends recent work to synthesize existing data and assess the status of NCC salmon by incorporating habitat-based priors for carrying capacity (Connors et al. 2018). While survival and abundance of sockeye have declined for some populations on the NCC in recent years (Peterman and Dorner 2012, Connors et al. 2016), we found that 32 of the 54 populations with recent escapement data are likely to be at or above spawner abundances that maximize yield (S_MSY_), however these escapement data extend only to 2017 and recent returns to many sockeye populations have been near the lowest on record (Fisheries and Oceans Canada 2019). These results demonstrate the promise of combing stock-recruit based population assessments with habitat information to reduce uncertainty in stock-recruit model estimates of population parameters that underpin management and recovery planning for sockeye and other freshwater-rearing salmon.

Given the dearth of stock-assessment for many NCC sockeye populations, there is an opportunity to link existing data on spawner and recruit abundance, and habitat-derived estimates of lake capacity to generate conservation benchmarks and management targets for data-poor populations across the region. Overall, models that shared information across populations and integrated lake-based estimates of carrying capacity reduced uncertainty in estimates of demographic parameters for our 70 populations of sockeye salmon. In recent decades, hierarchical models assuming a common distribution for productivity (*α*) have become commonplace in fisheries stock-assessment (Thorson and Minto 2015), as populations of the same species often face the same fundamental constraints on their productivity (Myers and Mertz 1998). In our analysis, hierarchical models produced more biologically-conservative estimates of per-capita productivity at low abundance (*α*), than single population models, and reduced uncertainty in estimates of productivity. At the population level, shrinkage in estimates of the productivity parameter between the population-specific and regional-hierarchical models were most pronounced in populations with limited data, or where the assumption of independent productivity values produced extreme estimates of α. Reduced uncertainty, and shrinkage towards the hierarchical mean (*mu*.α) are significant from a conservation and management standpoint because productivity has a direct influence on estimates of sustainable harvest rates, and the expected rate of population recovery from short-term downturns in population size. Fortunately, shrinkage in productivity (*α*) values towards the hierarchical mean produced only modest changes estimates of spawner abundance at MSY (S_MSY_). However, in general this shrinkage towards hierarchical mean productivity (*α*) will result in more biologically-conservative estimates of sustainable harvest rates in populations with high productivity but less biologically-conservative harvest management in populations with low productivity.

Inclusion of prior information on lake productivity or size also served to reduce uncertainty in estimates of spawner abundance at carrying capacity (S_MAX_). Given the fundamental constraint of lake size and productivity for most lake-rearing sockeye populations (Groot and Margolis 1991, Shortreed et al. 2001), inclusion of this information through the specification of priors is a logical way to narrow the range of possible values for S_MAX_ in a population. However, S_MAX___PR_ was often lower than stock-recruit based estimates of carrying capacity and model estimates of S_MAX_ tended to be lower when lake-PR priors were used, leading to lower estimates of S_MSY_. Perhaps these populations use additional habitats for rearing (e.g., estuary or river) or estimates of lake productivity could have changed through time. Alternatively, systems where S_MAX___PR_ is well above S_MAX_ may have population capacity set by spawning habitat, not lake rearing habitat. Management targets estimated with lake-PR priors should therefore be implemented with precaution, particularly when S_MAX___PR_ falls well below stock-recruit based estimates of capacity. Despite this call for caution, our estimates of S_MAX_ and S_MSY_ were consistent with previous escapement targets derived from a variety of stock-recruit and habitat-based approaches in well studied populations like Meziadin Lake (Bocking et al. 2002), suggesting that these approaches can add to and strengthen the scientific basis of management, reducing uncertainty in data-limited salmon fisheries.

The survival and productivity of sockeye populations in coastal British Columbia has fluctuated over time, and in recent decades smolt-to-adult survival is believed to have declined for many stocks (Peterman and Dorner 2012). Population size at carrying capacity (S_MAX_) and spawner escapements yielding optimum recruitment (S_MSY_) vary in relation to marine survival (Moussalli and Hilborn 1986; Atlas et al. 2015), and recent declines in survival mean that current S_MAX_ values are likely lower than they were historically when marine survival was higher. This temporal variability in stock-recruit dynamics also creates the possibility of autocorrelation among recruitment residuals and population size that can introduce negative bias in estimates spawner abundance at carrying capacity (1/β) and positive bias in productivity (*α*) (Walters 1987). This bias will be greatest for unproductive populations with temporal autocorrelation in stochastic natural mortality (Korman et al. 1995, Myers and Barrowman 1995). Ideally, stock-recruit models would be fit with temporal autocorrelation in residuals or with time-variant productivity to account for temporal trends in recruitment variation (e.g. Liermann et al. 2010). However, given the frequency of missing data for many populations in our study we opted to treat productivity as time invariant, modeling stock-recruit dynamics across the entire six decades of population monitoring data, reflecting both current and historic recruitment dynamics. While this approach is valuable for evaluating stock status relative to historic conditions and quantifying the benefits of using habitat-based priors in model specification, models which incorporate temporal variability in productivity should be used to generate management targets and inform harvest planning.

Despite recent downward trends in productivity for sockeye in British Columbia, about half (30 of 54 with escapement data) of sockeye populations with at least three years of escapement data since 2000 were above spawner abundance at maximum sustained yield (S_MSY_) (mean = 1.16 times). While declining productivity has resulted in elevated conservation concern, harvest rates have declined by roughly 50% across the NCC over the last 25 years (English et al. 2016, Walters et al. 2019). Contemporary harvest rates are estimated to range between 10-30% allowing many populations to maintain abundance above their estimated S_MSY_. While these reductions in harvest rates have allowed recent escapements for many populations to remain above S_MSY_ the need for conservation in many years has contributed to a loss of fishing opportunities, creating major hardships for subsistence and commercial fishers (Connors et al. 2019; Walters et al. 2019; Steel et al. *in review*). Other populations (16 of 70 evaluated) lacked recent escapement data required to evaluate conservation status, highlighting some of the challenges associated with evaluating population status in remote regions of coastal BC. Twenty-two of the 70 populations evaluated had recent escapement levels below their median estimate of S_MSY_. Among the populations assessed by both models to be below S_MSY_, both Owikeno and Atnarko were historically important for commercial and food, social and ceremonial fisheries, and the collapse of these populations has led to severe reductions in fisheries openings with impacts on salmon dependent communities along the central coast (McKinnell et al. 2001, Connors et al. 2019).

Our work demonstrates the utility of limnological studies from sockeye rearing lakes to inform conservation and management targets. PR-based estimates of capacity can typically be made with data from one or a few years (Hume et al. 1996), whereas producing reliable estimates of capacity with stock-recruit modeling requires decades of continuous monitoring of abundance. Managers should interpret PR model outputs, whether independently or as priors in stock-recruit modeling carefully, particularly if PR model estimates of capacity fall well above or below stock-recruit estimates of capacity. However, spawner-recruit data may also yield biased inference about population carrying capacity, particularly if models are fit to short timeseries or if survival has undergone directional change.

Cases where estimates of carrying capacity from the PR model and stock-recruit models diverge may provide important clues about the factors limiting population size and productivity. Since PR models capture only autotrophic energy pathways they may underestimate lake productivity supporting sockeye, particularly in lakes where microbial pathways comprise a high proportion of total production (Stockner and Shortreed 1989, Atlas et al. 2020). In other cases, zooplanktivorous competitors such as three-spine stickleback (*Gasterosteus aculeatus*) may limit food availability for sockeye and reduce system-specific carrying capacity relative to the overall primary productivity of a given lake (O’Neill and Hyatt 1987, Shortreed et al. 2001), for example in Alastair Lake and many other coastal lake systems. Efforts have been made by some investigators to account for competitor biomass in PR-based capacity estimates (Cox-Rogers et al. 2004), however given the broad geographic scope of our study, and the lack of data on fish community composition for most of the lakes, we did not incorporate competitor biomass. This omission likely inflated PR-based estimates of capacity for lakes with large populations of competitors. In addition, the model assumes that all sockeye juveniles are lake rearing, and that density-dependent population regulation occurs in lakes. While lake rearing is certainly the dominant juvenile life-history for sockeye on the BC coast, ocean-type and stream-rearing life histories have also been documented in the several areas of the NCC (e.g. Beveridge et al. 2015, Connors 2016) and undocumented life-history diversity likely exists in many other populations in the region. If stream-or ocean-rearing sockeye are included in spawner enumeration, their contribution to the population will inflate S_MAX_ relative to the productivity capacity of the rearing lake (S_MAX___PR_). Finally, population growth may be limited by the amount of available spawning habitat, rather than the size and productivity of the rearing lake (e.g. Bear Lake) (Shortreed et al. 1998). In these instances, the PR model will fail to capture the habitat processes that limit population growth, leading to higher estimates of S_MAX_ when PR-based capacity values are integrated with stock-recruit models as prior information. Regardless of these important biological considerations, our study highlights the utility of the PR model in representing the productive capacity of sockeye salmon nursery lakes, with strong coalescence in predicted carrying capacity from traditional stock-recruit and PR model approaches across a diversity of coastal sockeye stocks.

The integration of hierarchical-Bayesian stock-recruit models with prior information on lake-productivity is a promising avenue towards better informed management of sockeye in the NCC. However, understanding the limitations and potential biases associated with the dataset and modeling approaches are important if model outputs are to be used to guide management. For instance, enumeration methods are not uniform between study systems and may have changed through time, and observation error in escapement estimates likely varies across populations. While salmon populations tend to show regional coherence in population trends (Pyper et al. 2005), the absence of recent population monitoring data may produce inaccurate assessments of regional populations status if reductions in monitoring efforts have been biased towards underperforming populations (Price et al. 2008). Finally, for most populations – with the exception of Babine, Atnarko, Owikeno, Long, and Meziadin – only average age structure was available, and we assumed that age structure was fixed across brood years. Zabel and Levin (2002) have cautioned against the use of fixed age structure in stock-recruit models, as it will tend to smooth recruitment variability, resulting in underestimates of S_MAX_ and overestimates of productivity.

The model we developed can serve as a roadmap for future efforts to estimate conservation and management goals for data limited salmon stocks. While the demographic parameter estimates we present are preliminary, the modeling approach provides a foundation for conservation and fisheries management under the Wild Salmon Policy (DFO 2005) and the Fisheries Act. Our results suggest that despite reductions in productivity and abundance in recent decades, most sockeye populations on the NCC with intact habitats and moderate harvest rates are at or above MSY and presumably of low immediate conservation concern. Given these findings it is likely that populations currently above S_MSY_ may continue to support limited directed fisheries under precautionary management approaches.

Our analysis demonstrates the utility of merging hierarchical stock-recruit analysis with information derived from habitat-based models. Hierarchical models are a powerful tool for estimating stock-recruit parameters in populations with variable data quality or quantity, and the inclusion of habitat-based priors on carrying capacity provides a logical and biologically grounded means of defining priors for S_MAX_. Combining insights from habitat-based models of population capacity with stock-recruit analyses in a Bayesian framework can reduce uncertainty associated with estimates of population parameters and management targets resulting from highly stochastic adult-to-adult recruitment data. While we advise caution in setting management targets for fisheries with limited data, habitat-based models are a useful starting point for evaluating stock status and setting precautionary harvest goals even with limited stock-recruit data. Together these approaches provide key tools for managers of fisheries in developing economies or remote landscapes where a lack of population data currently hinders scientific decision making.

**Figure 6:**
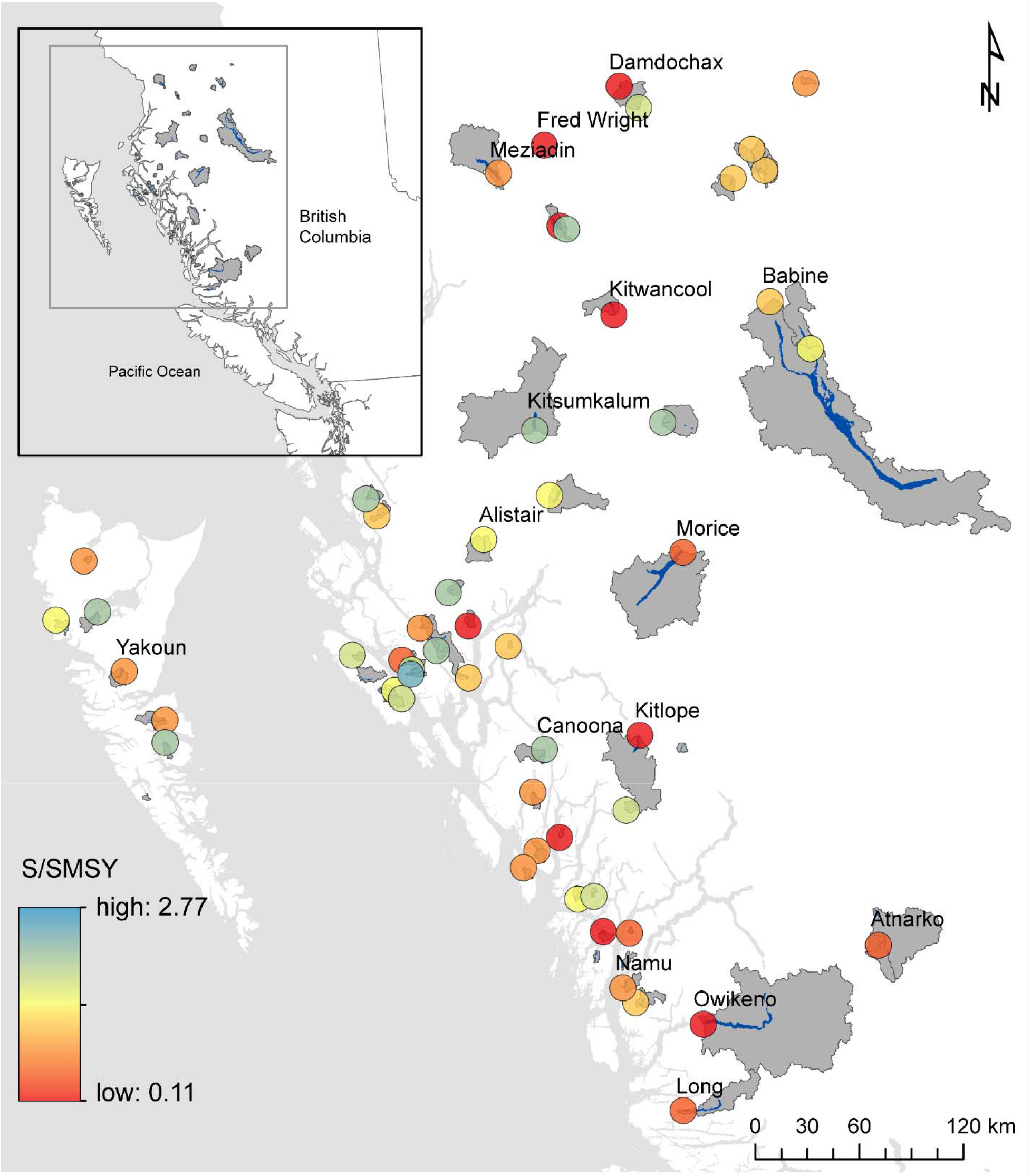
Estimated status (S/S_MSY_) for the 54 populations of sockeye on the NCC with recent escapement data.

## Acknowledgements

This work would not have been possible were it not for decades of work by countless researchers and technicians at DFO, with particular acknowledgement is due to Jeremy Hume (retired), John Stockner (retired), and the late Ken Shortreed, who developed the PR model and collected and analyzed limnological data from sockeye lakes for decades to improve our collective understanding. We are grateful for the efforts of myriad stock-assessment biologists and stream counters who gathered salmon escapement data. Will Atlas receives funding support through a Hakai Fellowship at Simon Fraser. Jonathan Moore is supported by the Liber Ero Foundation. We thank Thomas Buehrens for his advice on stock-recruit modeling, Mike Malick for code and input during early-stages of model development, and Kara Pitman for her help generating geospatial data and mapping sockeye lakes. We also thank Ryan Whitmore and the Gitxsan Watershed Authority for updated Slamgeesh weir data.

---

**Figure S1:**
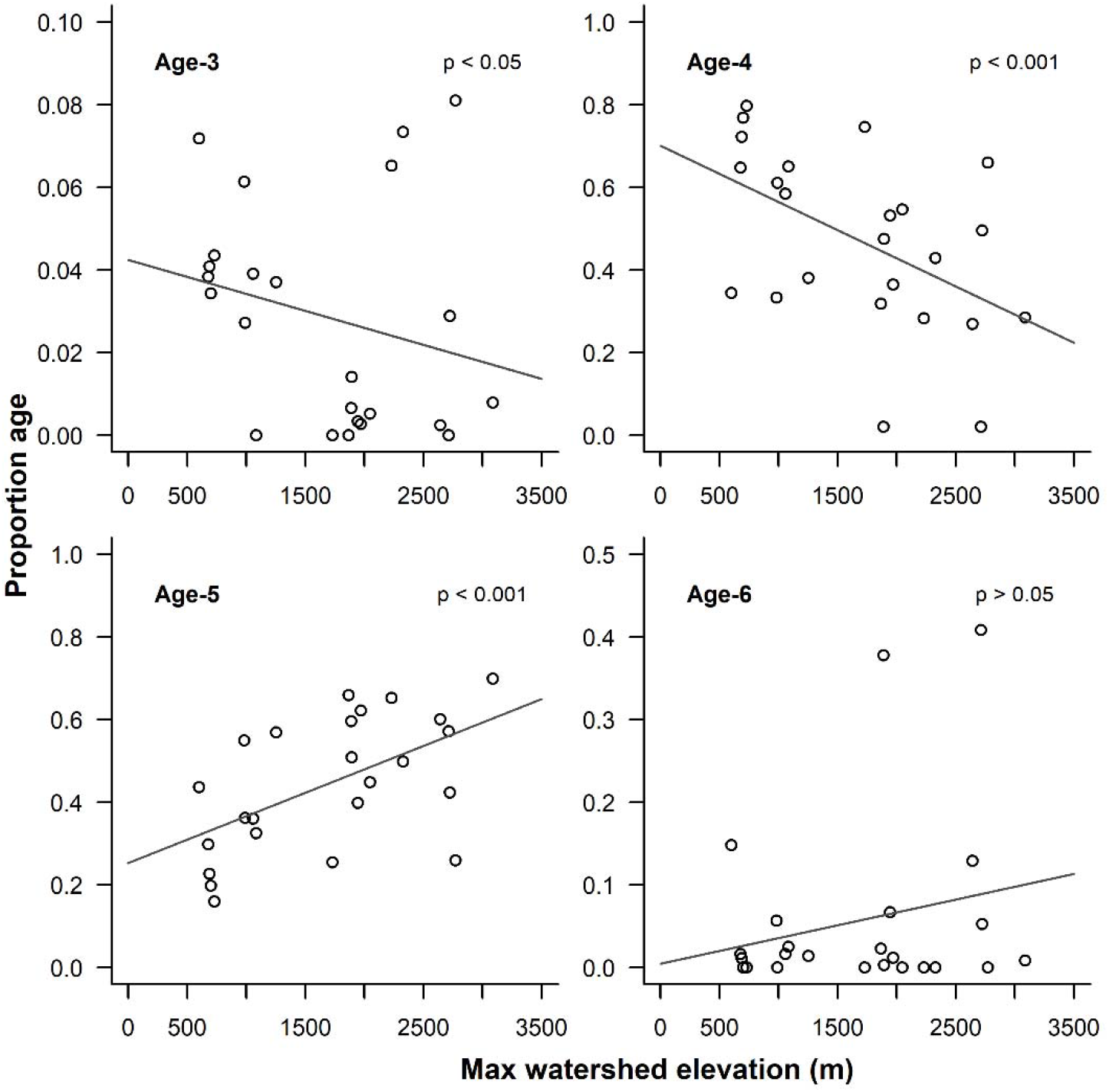
Relationship between mean population age composition and maximum watershed elevation (m), with p-values from multinomial regression fits reported in the top right corner of each panel.

